# The molecular basis of hypercontractility caused by the hypertrophic cardiomyopathy mutations R403Q and R663H

**DOI:** 10.1101/543413

**Authors:** Saswata S. Sarkar, Darshan V. Trivedi, Makenna M. Morck, Arjun S. Adhikari, Shaik N. Pasha, Kathleen M. Ruppel, James A. Spudich

## Abstract

Hypertrophic cardiomyopathy (HCM) mutations in ß-cardiac myosin and myosin binding protein-C (MyBP-C) cause hypercontractility of the heart. We show that hypercontractility caused by the HCM myosin mutation R663H cannot be explained by changes in the fundamental parameters such as actin-activated ATPase, intrinsic force, velocity of pure actin or regulated thin filaments, or the pCa50 of the velocity of regulated thin filaments. The same conclusion was made earlier for the HCM myosin mutation R403Q (Nag et al. 2015). Using enzymatic assays for the number of functionally-available heads in purified human ß-cardiac myosin preparations, we provide evidence that both R403Q and R663H HCM myosin mutations cause hypercontractility by increasing the number of functionally-accessible myosin heads. We also demonstrate that the myosin mutation R403Q, but not R663H, ablates the binding of myosin with the C0-C7 fragment of myosin binding protein-C.

## Introduction

The sarcomere of cardiac muscle is composed of actin-containing thin filaments and myosin-containing thick filaments. Sarcomeric contraction is driven by the cyclic interaction between actin and myosin in the presence of ATP. The resultant power generation is equated to the ensemble force (F_E_) generated by myosin heads on actin multiplied by the velocity (v) of actin filaments driven by myosin heads. Ensemble force is the product of the intrinsic force (F_i_) generated by a single myosin head on actin, the duty ratio (dr), which is the fraction of time a myosin molecule remains bound to actin, and the total number of functionally-accessible myosin heads (N_a_) for interaction with the actin filaments (*1*). Cardiac myosin has two N-terminal globular head domains (Subfragment 1, S1) and a long coiled-coil tail. The N-terminal ~40% of the tail is called Subfragment 2 (S2). The two S1 domains attached to the S2 domain constitute heavy meromyosin (HMM). The C-terminal ~60% of the tail is called light meromyosin (LMM). The LMM domain self assembles into thick filaments, which myosin binding protein-C (MyBP-C), titin and other sarcomeric proteins are associated with in the muscle sarcomere (*2–4*). MyBP-C is an elongated multidomain protein consisting of IgG- and Fn-like domains (*5*). There is approximately one MyBP-C molecule for every three myosin molecules in the C-zone of the myosin thick filament, or about one for every six myosin molecules when considering the entire thick filament (*4*, *6*).

The heads of myosin in cardiac muscle thick filaments can exist in at least two different states, an open state which is able to interact with actin and a closed state where the mesa domains (*7*) of the myosin heads are folded back onto their own S2 tail and are not accessible for interaction with actin. The best structural model of this folded-back sequestered state is the interacting heads motif (IHM) (*8–13*).

In relaxed muscle where the heads are unable to bind to the tropomyosin.troponin (Tm.Tn) regulated thin filaments, the open state heads are disordered (disordered relaxed state, DRX) and believed to be turning over ATP at a rate of ~ 0.03 s^−1^ (two orders of magnitude slower than the actin-activated rate of ~ 3 s^−1^), while the heads in the IHM state are believed to be in a super relaxed state (SRX) with an even lower ATP turnover of ~ 0.003 s^−1^ (*14*, *15*). Thus, SRX is defined as a very slow ATP turnover state of myosin as measured by a fluorescent-ATP assay (*14*), while IHM is a structural state in which the myosin heads fold on their own tail leading to an off state of the thick filament. The SRX and IHM states are correlated but there can be circumstances where a non-IHM state can also lead to SRX-like rates (*15*).

An attractive model is that the equilibrium of myosin between the IHM and the open state in cardiac fibers regulates the power generation of the sarcomere by controlling N_a_ (*7*, *16*). Phosphorylation of the regulatory light chain (RLC), one of the two light chain proteins that bind to myosin heads, is known to destabilize ordered myosin heads (*11*, *17*, *18*), leading to an increase in sarcomere contraction by increasing N_a_.

The C-terminal domains of MyBP-C (C8-C10) are associated with the thick filament LMM backbone (*19*, *20*). Biochemical studies have shown that the N-terminal domains (C0-C2) can interact with actin (*21*, *22*), the myosin RLC (*23*), the proximal part of S2 (proximal S2) (*24*), as well as short S1 (sS1, the myosin head fragment ending just after the ELC and therefore missing the RLC) (*11*). The interaction of C0-C2 with myosin has been postulated to sequester heads into a state not accessible for binding to actin, perhaps the IHM state, reducing N_a_ (*11*), while its interaction with the Tm.Tn-regulated thin filaments activates the regulated thin filaments at low Ca^2+^ concentration and can act as a load against contraction at high Ca^2+^ concentration (*25–28*).

Myosin and MyBP-C are the two major proteins involved in genetic hypertrophic cardiomyopathy (HCM), accounting together for ~70% of all known HCM mutations, ~35% in each (*29*). Hypertrophic cardiomyopathy (HCM) is a prevalent genetic cardiac disease affecting ~ 1 in 500 people (*30*). HCM results in hypercontractility of the heart, followed by hypertrophy, fibrosis, myocardial disarray, diastolic dysfunction and sometimes sudden death among HCM patients (*30*, *31*).

Here, we investigate two myosin HCM mutations, R403Q and R663H, which lie on the myosin mesa surface (*7*), using biochemical and biophysical assays as well as binding of MyBP-C. A previous study showed that the R403Q HCM mutation has minimal effects on the basic contractility parameters of the human ß-cardiac sS1, which are unlikely to account for the hypercontractility seen clinically (*32*). Here we show this is even more dramatically true for the R663H HCM mutation, which leads to virtually no changes in the fundamental contractile behaviors of human ß-cardiac myosin. Our recent work has focused on the ‘unifying hypothesis’ (*7*, *16*) that most HCM mutations may be releasing myosin heads from a sequestered state, thus increasing the number of heads functionally accessible for interaction with actin (N_a_). The IHM state is the likely structural analog of this sequestered state (*9*, *10*), which fits with the findings that the majority of HCM mutations are found on the myosin mesa, proximal S2, the converter domain, and other key intramolecular interaction sites in the IHM structure (*12*, *13*, *33*).

Here we show, using functional assays for myosin activity, that a major effect of both the R403Q and R663H mutations is to release heads from their sequestered state, which results in more heads becoming accessible for actin interaction (that is, increasing N_a_). In addition, we show that the R403Q mutation, but not the R663H mutation, dramatically reduces the binding of the C0-C7 fragment of MyBP-C to human ß-cardiac myosin. These results are discussed in relation to the locations of these two mutations in the model IHM structure.

## Results

### The HCM mutation R663H has no effect on fundamental parameters of myosin-based contractility

The R663H HCM mutation is nearly unique among the HCM mutations that have been studied at the molecular level in that it has no significant effect on the fundamental parameters of myosin-based contractility (Fig. 1, Table S1). The k_cat_ of the actin-activated ATPase activity of R663H human ß-cardiac sS1 (2.9 ± 0.1 s^−1^) is nearly identical to that of WT human ß-cardiac sS1 (WT, 3.0 ± 0.2 s^−1^) (Fig. 1A), and the intrinsic force of the mutant and WT forms are within 10% of one another (WT, 1.4 ± 0.1 pN; R663H, 1.5 ± 0.1 pN; Fig. 1B, Fig. S1). The velocities in the in vitro motility assay with both actin alone (WT, 820 ± 39 nm/s; R663H, 834 ± 31 nm/s) (Fig. 1C) and with regulated thin filaments at pCa = 4 (Fig. 1D) (WT, 1019 ± 25; R663H, 944 ± 42 nm/s) are not significantly different from the WT human ß-cardiac sS1. Additionally, the Ca^2+^ concentration dependence of regulated thin filament velocities of the WT (Fig. 1E) and R663H (Fig. 1F) human ß-cardiac sS1 constructs are nearly identical. Fitting the data with the Hill equation yielded the pCa_50_ values of 6.3 and 6.4 (obtained from the fitting of data averaged for 4 different experiments) for WT and R663H sS1, respectively.

**Figure 1.**
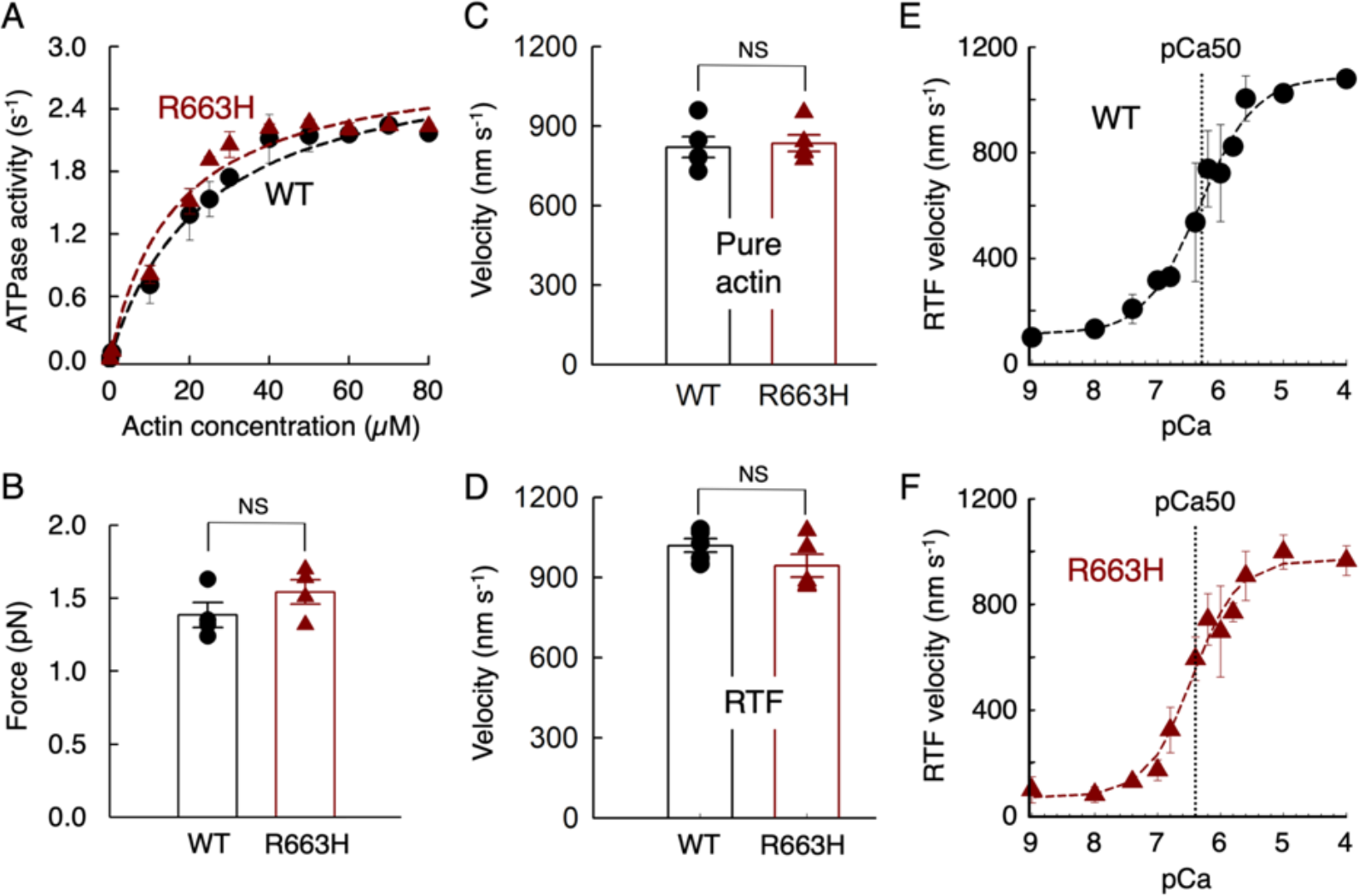
Fundamental contractile parameters of WT (black) and R663H (dark red) human β-cardiac sS1. (**A**) Actin-activated ATPase data for WT and R663H sS1. Fitting the traces yielded k_cat_ values of 3.0 ± 0.2 and 2.9 ± 0.1 s^−1^ for WT and R663H sS1, respectively. Dashed lines are the fitted lines. Shown is representative data, the average of two experiments from single preparations of both proteins. (**B**) Intrinsic force measurements of WT (black) and R663H (dark red) human β-cardiac sS1using an optical trap. The individual force values were averaged from four single molecule force measurements. (**C**) Comparison of in vitro motility of actin filaments for WT (black) and R663H (dark red) human β-cardiac sS1 (5 different experiments from 5 independent protein preparations for each of WT and R663H sS1). (**D**) Comparison of in vitro motility of regulated thin filaments at pCa = 4.0 for WT (black) and R663H (dark red) human β-cardiac sS1 (5 different experiments from 4 independent protein preparations for each of WT and R663H sS1). (**E**) Ca^2+^ sensitivity measurements for WT human β-cardiac sS1 (4 different experiments were averaged). (**F**) Ca^2+^ sensitivity measurements for R663H human β-cardiac sS1 (4 different experiments were averaged). The data for (E) and (F) were fitted (dashed lines) to the Hill equation to estimate the pCa50. For all panels, error bars denote SEM. NS, not significant, P > 0.05. (B) P = 0.24; (C) P = 0.78; (D) P = 0.18.

The IHM consists of a blocked head (the actin binding domain is blocked) and a free head (the actin binding domain is available). The blocked head interacts with proximal S2 and with the free head. The positions of residue Arg663 (light blue) on the IHM homology model of human ß-cardiac myosin constructed by Robert-Paganin et al. (*13*) are shown on the blocked head mesa (salmon) and free head mesa (pink) in Fig. 2A. The blocked head Arg663 (bh R663) is near but not interacting with the proximal S2, and the free head Arg663 (fh R663) is near the interface of a head-head interaction site (Fig. 2A,B). Given the closeness of Arg663 to the proximal S2 in the model, we measured the K_D_ for binding of proximal S2 to the WT (67 ± 12 μM; Fig. 2C) and R663H (50 ± 6 μM; Fig. 2D) human ß-cardiac sS1. They were not significantly different. This is consistent with the IHM model, where Arg663 is not in direct contact with proximal S2 (Fig. 2B). We previously demonstrated that R403Q human ß-cardiac sS1, which is also not in direct contact with proximal S2 in the IHM model, did not show a change in K_D_ for binding to proximal S2 (*11*). In contrast, Arg249, His251 and Arg453 are all directly underneath the proximal S2 in the IHM model (Fig. 2B) and the HCM mutations R249Q, H251N and R453C all significant weaken the binding affinity of human ß-cardiac sS1 to proximal S2 (*11*, *34*).

**Figure 2.**
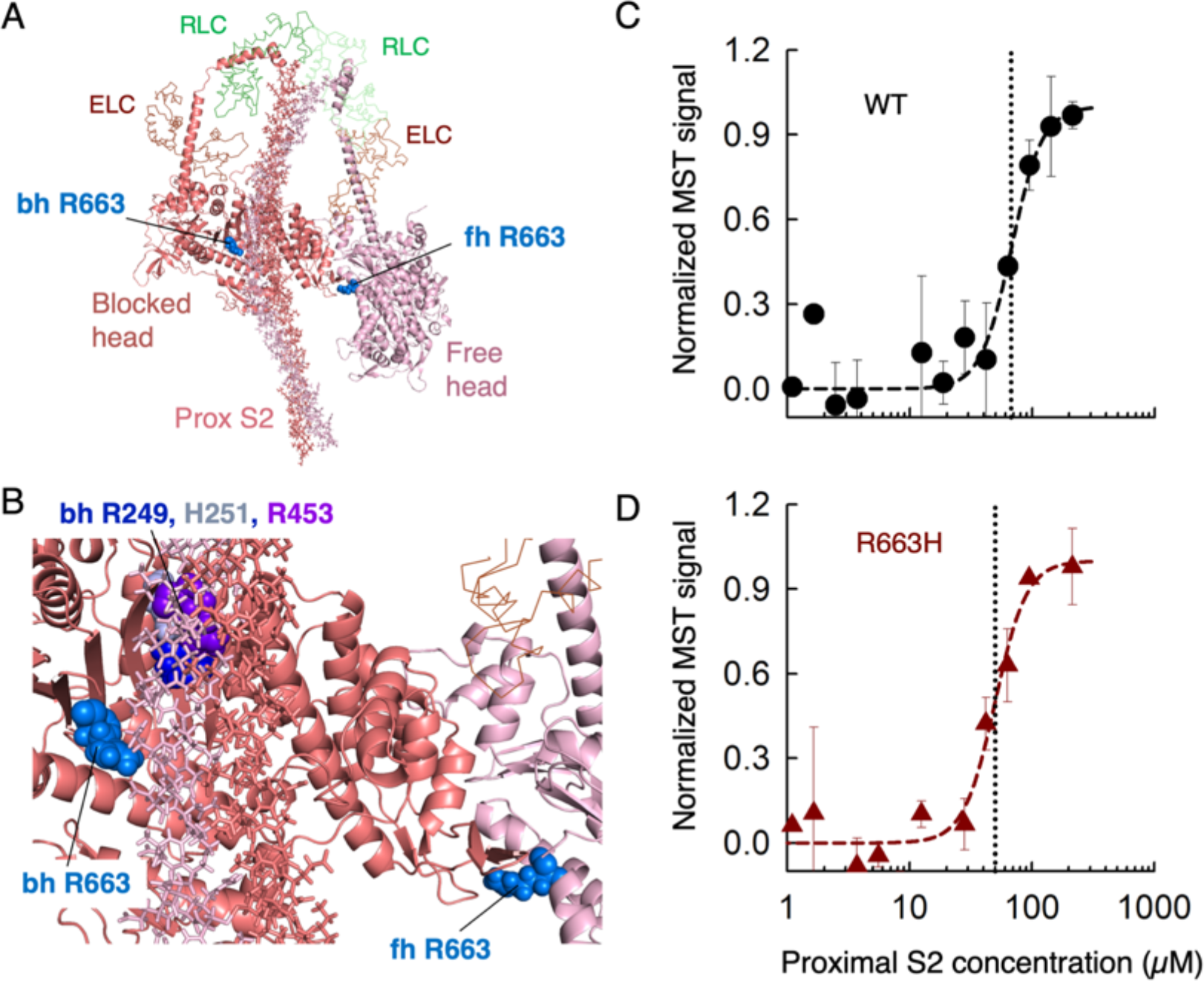
The affinity of WT and R663H human ß-cardiac sS1 binding to proximal S2. (**A**) Structural model of the IHM state for human β-cardiac myosin (Human sequestered state model; from Robert-Paganin et al. (*13*)). The S1 portion of the heavy chain are shown in cartoon mode (PyMol), the light chains in ribbon mode and the proximal S2 in stick mode. The positions of the blocked head (bh) and free head (fh) Arg 663 residues are shown as spheres in light blue. (**B**) A blowup of the IHM model showing the relationships between the Arg 663 residues and the proximal S2 position and the head-head interaction zone. Bh R249 (blue), H251 (grey) and R453 (purple) are shown for reference. (**C**) MST binding data for WT human ß-cardiac sS1 and proximal S2 in 100 mM KCl. (**D**) MST binding data for R663H human ß-cardiac sS1 and proximal S2 in 100 mM KCl. Data in C and D represent two measurements from a single set of protein preparations.

We conclude that the hypercontractility caused by the HCM mutation R663H, like R403Q (*32*), cannot be explained by changes in ATPase, velocity or intrinsic force, or the interaction between the S1 domain and proximal S2. We therefore explored whether an alternate mechanism could be that the R663H and R403Q mutations release myosin heads from their super relaxed state (SRX), which would provide more heads available for interaction with actin and thereby cause the hypercontractility seen clinically.

### The HCM mutations R403Q and R663H decrease the percentage of SRX in preparations of human ß-cardiac 25-hep HMM

One hallmark of the ability for a population of myosin molecules to form the IHM is a reduction in the apparent k_cat_ of the actin-activated ATPase activity of 25-hep HMM (which contains proximal S2; Fig. S2) compared to that of 2-hep HMM (which does not contain proximal S2; Fig. S2) {Adhikari, 2019 #1386}. Thus, WT human ß-cardiac 25-hep HMM (k_cat_ = 1.4 ± 0.1 s^−1^) has 58 ± 3% of the ATPase activity of WT 2-hep HMM (k_cat_ = 2.4 ± 0.1 s^−1^) (*35*) (Table S1). Both R403Q and R663H human ß-cardiac 2-hep HMM, however, have the same activity as the WT 2-hep HMM (R403Q, 2.7 ± 0.2 s^−1^; R663H, 2.8 ± 0.2 s^−1^) (Fig. 3A,B). Importantly, the 25-hep HMM versions of the R403Q and R663H human ß-cardiac myosins do not show the same ~40% decrease in activity compared to the 2-hep versions seen for the WT construct (Fig. 3A,B; Table S2). Both the R403Q and R663H human ß-cardiac 25-hep HMMs show higher activity (R403Q, 2.2 ± 0.2 s^−1^; R663H, 1.9 ± 0.1 s^−1^) than the WT 25-hep HMM (1.4 ± 0.1 s^−1^) (*35*), suggesting that these mutations cause heads to be released from the IHM state. These results suggest that there is a proximal S2-dependent stabilization of the IHM for WT human ß-cardiac 25-hep HMM that is destabilized by the presence of either the R403Q or the R663H mutation. We speculate that these effects may be a result of a weakening of the interaction between the blocked head with the free head in the IHM structure, where the bhR403 and fhR663 residues may play a role (Fig. 3G, H). We also hypothesize from these results that appropriate interaction of the blocked and free heads depends on the presence of proximal S2 to form the IHM structure.

**Figure 3.**
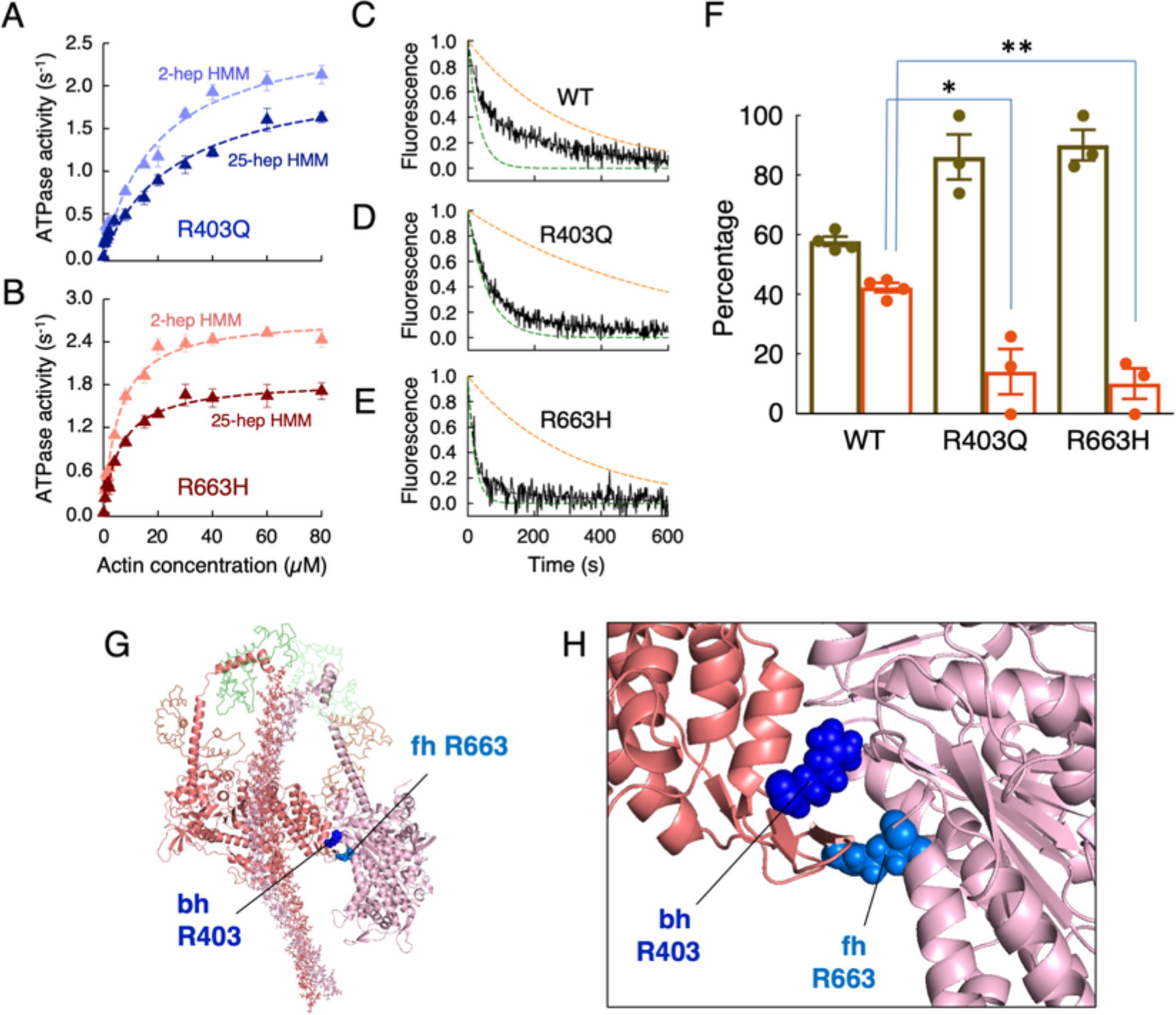
Functional assays for ATP turnover by WT, R403Q and R663H human ß-cardiac 25-hep HMM. (**A**) Actin-activated ATPase of R403Q 2-hep HMM (light blue) and R403Q 25-hep HMM (dark blue). (**B**) Actin-activated ATPase of R663H 2-hep HMM (orange) and R663H 25-hep HMM (dark red). For panels A and B, data is combined from 2 experiments from 1 protein preparation. Each point is an average with the error bar as SEM. (**C**) Fluorescence decay of mant-nucleotide release from WT human ß-cardiac 25-hep HMM in 25 mM KAc (solid black curve). Fitting the traces to a double-exponential equation yielded the DRX and SRX rates of 0.030 and 0.0034 s^−1^. The simulated single-exponential orange dashed curve is the SRX rate (0.0034s^−1^), and the simulated single-exponential dashed green curve is the DRX rate (0.030 s^−1^). The simulated orange and green dashed curves act as visual references for the single exponential fits of slow and fast phases, respectively, that derive from fitting the solid black data curves with the best two exponential fits. Thus, the data (black lines) were fit with a combination of these two single exponentials. (**D**) Fluorescence decay of mant-nucleotide release from R403Q human ß-cardiac 25-hep HMM in 25 mM KAc, where the fast and slow fitting rates were 0.017 s^−1^ and 0.0017 s^−1^ respectively. (**E**) Fluorescence decay of mant-nucleotide release from R663H human ß-cardiac 25-hep HMM in 25 mM KAc, where the fast and slow fitting rates were 0.047 s^−1^ and 0.0031 s^−1^ respectively. Panels C-E show representative data from 1 preparation of each protein. (**F**) Percentage of myosin heads in the SRX (orange) versus DRX (olive green) states calculated from the amplitudes of the double-exponential fits of the fluorescence decays corresponding to the mant-nucleotide release from the 25-hep HMMs shown in panels C-E. The data are from 4 measurements (each from individual protein preparations done on different days) of WT 25-hep HMM and 3 measurements of R403Q and R663H 25-hep HMM. Error bars denote SEM. *P ≤ 0.05; **P ≤ 0.01. WT vs R403Q P= 0.03; WT vs R663H P = 0.008. (**G**) Homology model of the IHM state of human β-cardiac 25-hep HMM (Human sequestered state model from Robert-Paganin et al. (*13*)) showing the positions of the blocked head (bh) Arg 403 residue (dark blue) and the free head (fh) Arg 663 residue (light blue) at a head-head interaction site. (**H**) A blowup of the IHM model showing the positions of the bh Arg 403 and fh Arg 663 residues at the interface between the blocked head (dark salmon) and the free head (pink).

The definition of the SRX is a population of myosin molecules with a reduced level of single turnover basal ATPase rate (*14*). Thus, in the absence of actin, myosin heads that are free to interact with actin (disordered relaxed state, or DRX) turnover ATP at ~0.03 s^−1^, whereas the SRX turns over ATP at only ~0.003 s^−1^. These rates are measured by loading a fluorescent analog of ATP (mant-ATP) onto the myosin which results in a higher fluorescence, and then chasing with an excess of unlabeled ATP and watching the decrease in fluorescence as the mant-nucleotide dissociates from the myosin (Fig. 3C, WT myosin, black curve; Table S3). The decay rate shown in Fig. 3C cannot be fit by a single exponential, indicating a mixture of DRX and SRX turnover rates in the WT human ß-cardiac 25-hep HMM population. The amplitudes provide a measure of the percentages of DRX (58 ± 2%) and SRX (42 ± 2%) in the population at a particular ionic strength (Fig. 3F). In contrast to the WT myosin, R403Q and R663H human ß-cardiac 25-hep HMM showed only 14 ± 7% and 10 ± 5% SRX in the population, respectively (Fig. 3D-F; Table S3).

We have shown previously that SRX is correlated with formation of a folded-back sequestered state (*15*), probably the IHM. Thus, the decreased SRX that arises from these mutations suggests that they prevent formation of the IHM, in agreement with the actin-activated ATPase measurements. Taken together, these data support our hypothesis that bhR403 and fhR663 residues may play a role in stabilizing the IHM structure, and that the HCM mutations at these residues weaken the blocked head interaction with the free head (Fig. 3G,H), providing more myosin heads for interaction with actin.

### R403Q but not R663H disrupts the interaction of MyBP-C with human ß-cardiac 25-hep HMM

We and others have hypothesized that the N-terminal C0-C2 fragment of MyBP-C binds to a folded state of myosin (*11*). Previous binding data showed that both full length MyBP-C as well as C0-C2 bind to WT sS1 in a phosphorylation-dependent manner (*11*). Here, we examined the binding of the C0-C7 fragment of MyBP-C to human ß-cardiac 25-hep HMM using microscale thermophoresis (MST). We used this longer C0-C7 fragment in order to account for any and all effects that the myosin mutations may have in changing MyBP-C binding. We have left off the C8-C10 region, as it is known to interact with LMM (*19*, *20*).

The K_D_ for binding of R663H 25-hep HMM to C0-C7, following the HMM eGFP fluorescence and titrating with C0-C7 was 900 nM, nearly identical to that of WT 25-hep HMM (Fig. 4A). Similar results were obtained when we titrated fluorescently-labeled C0-C7 with 25-hep HMM (Fig. S3). In contrast, the R403Q 25-hep HMM showed no binding to C0-C7 up to 50 μM C0-C7 (Fig. 4A). One highly speculative model that fits all the previously known data for binding of C0-C7 to the IHM state of human ß-cardiac myosin is shown in Fig. 4C (only C0-C2 is shown since the position of C3-C7 is uncertain). The Arg403 residue on the blocked head of the IHM may interact with the M domain, and the Arg403 residue on the free head may interact with the C2 domain.

**Figure 4.**
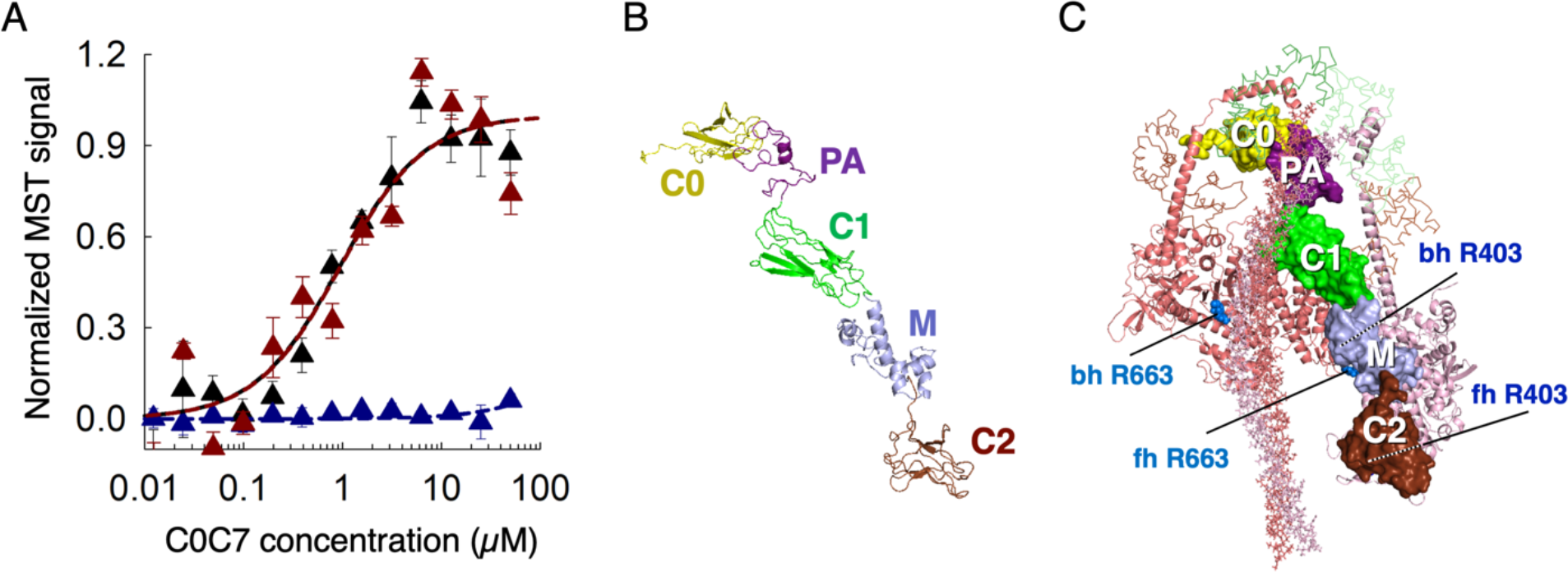
Binding data and model of human β-cardiac HMM with N-terminal MyBP-C. (**A**) Binding of human ß-cardiac 25-hep HMM with the C0-C7 fragment of MyBP-C for WT (black), R663H (dark red) and R403Q (blue). Data points represent the mean at each C0-C7 concentration with the s.e.m. shown. These representative data are the average of 3 measurements from a single set of protein preparations. (**B**) Homology model showing the N-terminal residues C0 to C2 of cardiac MyBP-C. (**C**) Backside view of a structural model of the IHM state for human β-cardiac myosin (Human sequestered state model from Robert-Paganin et al. (*13*)) with C0-C2 bound in the groove between proximal S2 and the mesa of the free head.

## Discussion

Both the R403Q (30) and R663H mutations have marginal effects on fundamental parameters of the actin-activated chemo-mechanical cycle. A previous study, however, showed that the R403Q and R663H mutations destabilize the SRX in functional studies of cardiac fibers, consistent with causing an increase in N_a_ (*15*). The results we present here complement and build on that work by showing that the R403Q and R663H mutations disrupt the SRX of purified human ß-cardiac 25-hep HMM, again consistent with their causing an increase in N_a_. Similar results were recently reported for the myosin HCM mutations R249Q, H251N, D382Y and R719W (*35*). These data all support a unifying hypothesis that the primary effect of most HCM mutations may be to increase the number of functionally available heads in the sarcomere, leading to the hypercontractility seen clinically. Furthermore, the locations of all these mutations are consistent with the IHM structural model for the folded-back sequestered state of human ß-cardiac myosin. The residue Arg403 is located at a particularly interesting location in the IHM model structure. It sits at a junction of three surface domains: the actin binding domain, the mesa and the primary head-head interaction site (PHHIS) (*16*), the latter two of which are important for IHM formation. The mutation does cause a 3-5 fold increase in the Km for actin binding (Table S2), but all other actin interaction parameters studied using sS1 show very little change by the R403Q mutation (*32*). The location of the R403Q mutation, however, may be expected to affect the stability of the IHM. This is consistent with what we report here. The free head R663H mutation similarly lies at a head-head interaction site within the IHM, and we propose that it therefore weakens the IHM complex.

The possible role of MyBP-C in sequestering heads in an off state is a pivotal area of future research, and here too the Arg403 residue seems to play an important role. The head domain of myosin showed a phosphorylation-dependent binding of the N-terminal domains of MyBP-C (*11*), and here we demonstrate binding of C0-C7 to the purified double-headed human β-cardiac myosin construct 25-hep HMM. The R403Q mutation essentially eliminates this binding. In our model of a C0-C2-HMM complex (Fig. 4C), both the free and blocked head Arg403 residues are potentially stabilizing the complex, and it therefore might not be surprising that the R403Q mutation so dramatically disrupts the complex. It must be emphasized, however, that this is a highly hypothetical model, and the structure of the MyBP-C-HMM complex is entirely unknown. While we have depicted the C0-C2-HMM complex as an IHM complex, it is entirely possible that MyBP-C is sequestering myosin heads away from interaction with actin by binding the heads in a different folded conformation or even in a more open configuration (*36*). Thus, a high priority in the field is to obtain even a low-resolution structure of the MyBP-C-HMM complex.

The potential importance of a folded-back sequestered state of cardiac myosin has recently been highlighted in diseases like HCM, and also in the mechanism of action of cardiac inhibitors like mavacamten (*15*, *37*, *38*). The cardiac activator omecamtiv mecarbil has also been shown to stabilize the open state of cardiac myosin while also impacting the kinetics of actin-bound myosin motors (*39–41*). Thus, further research into the determinants of the sequestered state of myosin and the role of MyBP-C in holding myosin heads in a reserved state can lead to novel insights into drug discovery for cardiac activators and inhibitors.

## Acknowledgements

The authors thank the Spudich laboratory members for their comments on the paper. This work was funded by NIH grants GM33289 and HL117138 (J.A.S.). D.V.T is supported by a Stanford Lucile Packard CHRI Postdoctoral Award (UL1 TR001085) and an American Heart Association Postdoctoral Fellowship (17POST33411070). M.M.M. was supported by the Stanford Cellular and Molecular Biology training grant. A.S.A was support by a Stanford Lucile Packard CHRI Postdoctoral Award (UL1 TR001085), Stanford CVI Postdoctoral Award, Stanford ChEM-H Postdocs at the Interface Award, and American Heart Association Postdoctoral Fellowship (16POST30890005). Author Contributions: S.S.S. and K.M.R. purified human ß-cardiac myosins. D.V.T. and M.M.M. purified MyBP-C. S.S.S. performed the experiments for measuring basic contractile parameters of sS1. S.S.S., D.V.T. and M.M.M. performed single ATP turnover and MST binding experiments with purified human ß-cardiac myosins and analyzed the data. A.S.A. performed actin-activated ATPase assays and the analysis of data. S.N.P. constructed the homology model of the C0-C2 domain of MyBP-C. S.S.S. and J.A.S. wrote the paper. S.S.S., D.V.T., M.M.M., A.S.A., S.N.P., K.M.R. and J.A.S. contributed to data analysis/interpretation and editing of the manuscript. Competing interests: J.A.S. is a co-founder of Cytokinetics and MyoKardia, biotechnology companies developing small molecules that target the sarcomere for a variety of diseases, and a member of their scientific advisory boards. K.M.R. is a member of the MyoKardia scientific advisory board.

## Materials and Methods

### Homology modeling of the human cardiac C0-C2 fragment of MyBP-C

#### Template Identification

The complete MyBP-C protein sequence, which consists of 1274 amino acids and has a calculated molecular weight of 140.78 kDa, was retrieved from the UniProtKB database (http://www.uniprot.org/) (accession number Q14896). Searching the RCSB Protein Data Bank (http://www.rcsb.org/) confirmed that the complete tertiary structure of MyBP-C was not publicly available (*42*). As the initial results from BLASTP with poor homolog templates and subsequently low optimal structure from prediction servers led us to fragment the sequence keeping functional regions intact based on SMART (*43*) and PFAM (*44*) domain definitions. The fragmentation of the MyBP-C sequence based on functional regions showed some optimal identical structural templates for C0, C1, M-domain and C2 regions from PDB (*42*).

#### Multi-template homology modeling of the N-terminus of MyBP-C

The templates for these protein fragments had a good similarity with existing proteins whose structures were known. The multi-template modeling of the N-terminus C0-C2 was carried using the Modeller tool (Modeller V9.20) (*45*). The template structures of the fragments and the alignments were used for multi-template homology modeling to generate an initial model. The modelled structures were energy-minimized to convergence using SYBYL software and then finally passed through the validation process using PROSA (*46*) and Procheck (*47*).

### Expression and purification of human β-cardiac myosin constructs

A modified AdEasy™ Vector System (Qbiogene, Inc) was used to generate WT and mutant (R403Q and R663H) human β-cardiac constructs as described in detail elsewhere (*11*, *48*, *49*). Three different truncated versions of myosin were created: sS1, 2-hep HMM and 25-hep HMM, consisting of residues 1-808, 1-855 and 1-1016, respectively.

The cDNA of two different truncated versions of MYH7 (residues 1-808), corresponding to a short S1 (sS1), followed by a flexible GSG (Gly-Ser-Gly) linker were made: a) a carboxy-terminal eGFP linker or b) a carboxy-terminal 8-residue (RGSIDTWV) PDZ-binding peptide.

The 2-hep cDNA consists of residue 1 to 855 of MYH7 followed by a GCN4 leucine zipper to ensure dimerization. The GCN4 is followed by an eGFP moiety flanked by flexible GSG linkers, and a carboxy-terminal 8-residue (RGSIDTWV) PDZ binding peptide. The 25-hep HMM construct is similar to the 2-hep HMM construct, except it has 25 heptad repeats (175 amino acids) of the S2 region (1-1016 residues). Human ventricular essential light chain (ELC) with an N-terminal FLAG tag (DYKDDDDK) and TEV protease site was co-expressed with the heavy chain using an adenoviral vector/mouse myoblast C2C12 system. Expression and purification of all three constructs were done as described previously (*11*).

### Purification of C0-C7

A C0-C7 construct with a C-terminal His-tag was codon optimized for expression in E. coli and was expressed using the pET-21 a (+) expression vector. Cells containing the C0-C7 plasmid were grown in Rosetta (DE3) pLysS cells, induced, and harvested as described by the manufacturer (Qiagen, Germany). The cells were then lysed in a lysis buffer containing 10 mM Imidazole, 50 mM NaH_2_PO_4_, 300 mM NaCl, 1 mM BME, 0.01 mg/mL leupeptin, 1 mM PMSF, and Roche cOmplete protease inhibitor cocktail. Lysis was carried out on an emulsiflex or a sonicator, and the lysate was clarified by centrifugation at 35,000 g for 35 min. The supernatant was loaded on a Ni-NTA column (GE) on an FPLC. The column was washed with 10 mM and 15 mM Imidazole followed by a linear elution gradient of 15 - 400 mM Imidazole. After analyzing the fractions on SDS PAGE, pure fractions were pooled together and concentrated using an Amicon 10kDa MWCO filter. These concentrated fractions were then diluted into a buffer containing 10 mM Imidazole pH 7.5, 4 mM MgCl_2_, and 1 mM DTT and loaded on an ion exchange Q-column (GE). The dilution step was essential to bring down the ionic strength to 60 – 70 mM for efficient attachment of the C0-C7 to the Q-column. The protein was eluted by a 0 – 100 % linear NaCl gradient and the peak fractions were analyzed by SDS-PAGE. Pure fractions were pooled, concentrated and finally loaded on a preparatory grade Superdex S200 (16/600) size exclusion column. The C0-C7 eluted as a peak from the SEC column with minor shoulder peaks. Only the major peak fractions were collected, analyzed by SDS-PAGE, pooled and subsequently used for MST experiments. The protein was eluted in a buffer containing 10 mM Imidazole pH 7.5, 100 mM potassium acetate, 4 mM MgCl_2_, 1 mM EDTA and 1 mM DTT.

### In vitro motility assay

The motility assay was performed as described in detail previously (*49*, *50*). Both WT and mutant protein(s) were purified simultaneously and studied on the same day to minimize variability. All the experiments were performed at 23°C. Glass coverslips (VWR micro cover glass) were coated with a mixture of 0.2% nitrocellulose (Ernest Fullam Inc.) and 0.2% collodion (Electron Microscopy Sciences) dissolved in amyl acetate (Sigma) and air-dried before use. A permanent double-sided tape (Scotch) was used to construct four channels in each slide, and four different experiments were performed on the same slide. Partially inactivated myosin heads in S1 preparations were removed by the ‘dead-heading’ process before performing the motility assay. The process of ‘dead-heading’ had the following steps: A ten-fold molar excess of F-actin was added to myosin in the presence of 2 mM ATP; the mixture was incubated for 15 min in an ice bucket; 50 mM MgCl_2_ was added to form F-actin Mg^2+^-paracrystals and incubated for 5 min; the paracrystals were sedimented at 350,000g for 15 min; the supernatant was collected, and the sS1concentration was measured using the Bradford reagent (Bio-Rad). Before any experiments, dead-headed sS1 was diluted in 10% ABBSA [assay buffer (AB; 25 mM imidazole, pH 7.5, 25 mM KCl, 4 mM MgCl_2_, 1 mM EGTA, and 1 mM DTT) with bovine serum albumin (BSA, 0.1 mg/ml) diluted in AB], unless otherwise stated.

For motility experiments using pure actin, reagents were sequentially flowed through the channels in the following order: 10 μl of 4 μM SNAP-PDZ18 diluted in AB and incubated for 3 min; 20 ul of ABBSA to block the surface from nonspecific attachments and incubated for 2 min; 10 μl of a mixture of eight-residue (RGSIDTWV)-tagged human ß-cardiac sS1 (~0.05 to 0.1 mg/ml) and incubated for 3 min; 20 μl of AB to wash any unattached proteins; and finally, 10 μl of the GO solution [5 to 10 nM tetramethylrhodamine (TMR)-phalloidin (Invitrogen)-labeled bovine actin, 2 mM ATP (Calbiochem), an oxygen-scavenging system consisting of 0.2% glucose, glucose oxidase (0.11 mg/ml; Calbiochem), and catalase (0.018 mg/ml; Calbiochem)], and an ATP regeneration system consisting of 1 mM phosphocreatine (Calbiochem) and creatine phosphokinase (0.1 mg/ml; Calbiochem) in ABBSA].

For RTF motility experiments, reagents were sequentially flowed through the channels in the following order: 10 μl of 4 μM SNAP-PDZ18 diluted in AB and incubated for 3 min; 20 μl of ABBSA to block the surface from nonspecific attachments and incubated for 2 min; 10 μl of a mixture of eight-residue (RGSIDTWV)-tagged human ß-cardiac sS1 (~0.05 to 0.1 mg/ml) and incubated for 3 min; 10 μl of AB to wash any unattached proteins; 10 μl 5-10 nM (TMR)-phalloidin (Invitrogen)-labeled bovine actin and incubated for 3 min; 10 μl AB buffer to wash; 10 μl 400 nM tropomyosin.troponin complex and incubated for 6 min; 10 μl of the GO solution [2 mM ATP (Calbiochem), an oxygen-scavenging system (0.2% glucose, glucose oxidase (0.11 mg/ml; Calbiochem), and catalase (0.018 mg/ml; Calbiochem)), and an ATP regeneration system (1 mM phosphocreatine (Calbiochem) and creatine phosphokinase (0.1 mg/ml; Calbiochem))] in ABBSA.

For all experiments, movies were obtained using a Nikon Ti-E inverted microscope with an AndoriXon+EMCCD camera model DU885. All experiments were repeated with at least 4 different fresh protein preparations. At each condition, at least 3 different movies with duration of 30 s were recorded. Filament tracking and analysis of movies were performed using the previously published FAST (Fast Automated Spud Trekker) method, as described earlier (*50*).

### Single molecule force measurements

The experimental set up of the dual-beam optical trap is described in detail elsewhere (*51*). All experiments were performed at 23°C. Dead-heads of purified myosins were eliminated as described above. The eGFP tag at the C-terminal end of the myosin was used for surface attachment through binding with anti-GFP antibody. The nitrocellulose-coated glass surface of the sample chamber was also coated with 1.5 μm-diameter silica beads that acted as platforms. The steps followed for preparing the chamber were: 20 μl of anti-GFP antibody (~0.01 mg/ml) (Abcam) was flowed through the chamber; 20 μl of AB buffer containing 1 mg/ml BSA (ABBSA buffer) was flowed through to block the exposed surface; the surface was sparsely coated with myosin by flowing through ~50-200 pM of human β-cardiac sS1; the chamber was washed with 20 μl of AB buffer; 20 μl of ABBSA buffer containing 250-500 pM ATP, TMR-phalloidin–labeled biotin-actin filaments, neutravidin-coated polystyrene beads (Polysciences), and the oxygen-scavenging and ATP regeneration systems described above was flowed through the chamber; the chamber was sealed with vacuum grease to stop evaporation of the solution. Two neutravidin-coated polystyrene beads (1-μm diameter) were trapped in two different laser beams. Bead-actin-bead assembly, also known as a dumbbell, was formed using the trapped beads bound to each end of a TMR-phalloidin- and biotin-labeled actin filament. The dumbbell was stretched to remove compliance in the actin filament and brought close to the bead pedestal on the surface for interaction with myosin. A trap stiffness of ~0.1 pN/nm was used.

The data collected from individual molecules were analyzed for force measurements, and such experiments were performed using myosins from at least two protein purifications. The number of total force events from individual molecules on average was 20 to 300. We have always observed that the force distribution of all myosin constructs is accompanied with a long tail, as reported previously. Each force distribution was fitted to a double-Gaussian function to take account of the smaller population of higher force events in the analysis. The major peak of the fit yielded the intrinsic force of an individual molecule reported here. Such intrinsic force values of multiple molecules were used to calculate the mean force value (Fig. 1B).

### MST measurements

A detailed method of Microscale Thermophoresis (MST) for determination of the dissociation constant for the binding of two proteins was described previously (*11*). For all interactions, the unlabeled protein partner was titrated into a fixed concentration of the fluorescently-labeled partner (42 nM). Binding experiments between sS1 and proximal S2 were performed as described before (*11*). We carried out binding experiments between 25-hep HMM and C0-C7 by two methods. In one case C0-C7 was titrated against 42 nM 25-hep myosin, and in the other 25-hep HMM was titrated against 42 nM fluorescently labeled C0-C7. Sixteen such serially diluted concentrations of the unlabeled protein partner were prepared to generate one full binding isotherm. 25-hep and C0-C7 binding reactions were carried out in a buffer containing 10 mM Imidazole pH 7.5, 4 mM MgCl_2_, 1 mM EDTA, 1 mM DTT, 50 mM KAc, 1mM ATP and 0.05% Tween 20.

Samples were loaded into NT.115 premium treated capillaries (Nanotemper Technologies) after the reaction was incubated in the dark at 23°C for 30 min. The samples were then mounted in the Monolith NT.115 apparatus (Nanotemper Technologies) for binding measurements. All the data were recorded at 23°C. When myosin was held constant and C0-C7 was titrated, the myosin’s GFP fluorescence was measured by a blue LED at 30% excitation power (BLUE filter; excitation 460-480 nm, emission 515-530 nm) and IR-Laser power at 60% was used. C0-C7 was fluorescently labeled with a commercial His-tag labeling kit according to manufacturer instructions (Nanotemper technologies). His-labeled C0-C7 fluorescence was measured by a red LED at 60-90% excitation power (RED filter; excitation 605-645 nm, emission 680-685 nm) and IR-Laser power at 60% or 80% was used. Data analysis was performed with software NTAffinityAnalysis (Nanotemper Technologies) where the binding isotherms were derived from the raw fluorescence data. At least 2 independent measurements each from at least 2 different preparations of protein were carried out in each case. Representative binding curves are shown in Fig. 2 and 4.

### Actin-activated ATPase assay

All the experiments were performed using freshly prepared myosin at 23 °C as described previously (*11*). WT and R663H sS1, 2-hep HMM and 25-hep HMM mutant proteins were prepared simultaneously to minimize the effect of preparation variability. Myosin concentration was measured using eGFP absorbance. G-Actin was prepared as described previously (*11*). F-actin free of ATP was prepared by extensively dialyzing G-actin into ATPase buffer to remove any residual ATP. Actin concentration was then measured using absorbance at 290 nm in a spectrophotometer. This eliminates any contribution from the nucleotide at 280 nm.

The steady-state actin-activated ATPase activities of the human β-cardiac myosin preparations were determined using a colorimetric assay to measure inorganic phosphate production at various time points (0-30 min) from a solution containing myosin (0.01 mg/ml), ATP and increasing amounts of actin filaments (0-100 μM), ATPase buffer (10 mM imidazole pH7.5, 5 mM KCl, 1 mM DTT and 3 mM MgCl_2_). The time-dependent rate for each actin concentration was calculated by fitting the phosphate signal as a function of time to a linear function. The slope was then converted to activity units normalized to a single myosin head. The k_cat_ was extracted from the data by fitting the activity at each actin concentration to the Michaelis-Menten equation to determine maximal activity using the curve fitting toolbox in MatLab. The errors in the fitted values were determined using 100 bootstrap iterations.

### Single turnover experiments

Single turnover experiments and the analysis of data were described in detail previously (*15*). A plate-based fluorescence instrument (Tecan model – Infinite M200 PRO) was used to collect fluorescence data. Single turnover experiments were performed in a 96-well plate (Greiner polypropylene microplate). Experiments were performed with 3 different human ß-cardiac myosin constructs – WT, R403Q and R663H 25-hep HMM. 200 nM myosin in a buffer containing 10 mM Tris pH 7.5, 4 mM MgCl2, 1 mM EDTA, 1 mM DTT and 25 mM KAc was mixed with 2’-(or-3’)-O-(N-Methylanthraniloyl) adenosine 5’-triphosphate (mant-ATP, Thermo Fischer Scientific) at a final concentration of 200 nM. After 10 s, 4 mM ATP was added, followed by measuring the fluorescence signal at 470 nm after excitation at 405 nm for every ~2 s for 16 min. The fluorescence signals at time = 0 s and time = infinite were used to normalize the kinetic traces. Separate experiments were performed to get the fluorescence signal at time = 0 s by mixing 200 nM myosin and 200 nM mant-ATP and collecting the fluorescence signal. The fluorescence signal at time = infinite was obtained from the fluorescence measurement of 200 nM mant-ADP. The kinetic traces fitted to a bi-exponential decay function yielded the amplitudes and rates of fast (DRX rate ~0.03 s^−1^) and slow (SRX rate ~0.003 s^−1^) phases. The rates are reported with the mean and its standard error in Fig. 3F. All experiments were performed with at least 3 fresh myosin purifications for each construct.

