## Supplemental Materials for "The molecular basis of hypercontractility caused by the hypertrophic cardiomyopathy mutations R403Q and R663H"

| HCM mutation | Intrinsic force mutant/WT ratio | Velocity mutant/WT ratio | ATPase mutant/WT ratio |
|---|---|---|---|
| R403Q | 0.8 ± 0.1 | 1.2 ± 0.1 | 1.2 ± 0.1 |
| R663H | 1.1 ± 0.1 | 1.02 ± 0.02 | 0.9 ± 0.1 |

Table S1. Summary of mechano-chemical properties of human ß-cardiac R663H sS1 and R403Q sS1. The ratios of mutant/WT for basic mechano-chemical properties of sS1 are presented. The data for human ß-cardiac R403Q sS1 is taken from our published data (*32*).

| | 2-hep HMM $k_{cat}$ ($s^{-1}$) | 2-hep HMM $K_M$ (μM) | 25-hep HMM $k_{cat}$ ($s^{-1}$) | 25-hep HMM $K_M$ (μM) | 25-hep HMM: 2-hep HMM (%) |
|---|---|---|---|---|---|
| WT | 2.4 ± 0.1 | 4 ± 1 | 1.4 ± 0.1 | 9 ± 2 | 57 ± 3 |
| R403Q | 2.7 ± 0.2 | 20 ± 3 | 2.2 ± 0.2 | 27 ± 6 | 20 ± 8 |
| R663H | 2.8 ± 0.2 | 6 ± 1 | 1.9 ± 0.1 | 6 ± 2 | 33 ± 3 |

Table S2. Actin-activated ATPase activity values for human ß-cardiac 2-hep HMM and 25-hep HMM. The values of $k_{cat}$ and $K_M$ were obtained from the fitting of data shown in Fig. 3A (R403Q) and 3B (R663H). WT values were obtained from our previous work (*35*).

| | SRX rate (1/s) | % of SRX | DRX rate (1/s) | % of DRX |
|---|---|---|---|---|
| WT | 0.0037 ± 0.0004 | 42 ± 2 | 0.024 ± 0.003 | 58 ± 2 |
| R403Q | 0.0016 ± 0.0001 | 14 ± 8 | 0.013 ± 0.002 | 86 ± 8 |
| R663H | 0.0038 ± 0.0006 | 10 ± 5 | 0.032 ± 0.007 | 90 ± 5 |

Table S3. Summary of mant-ATP single turnover data. The mean SRX and DRX rates and the mean percentages of myosin heads in SRX and DRX are reported with SEM for human ß-cardiac WT, R663H and R403Q 25-hep HMM.

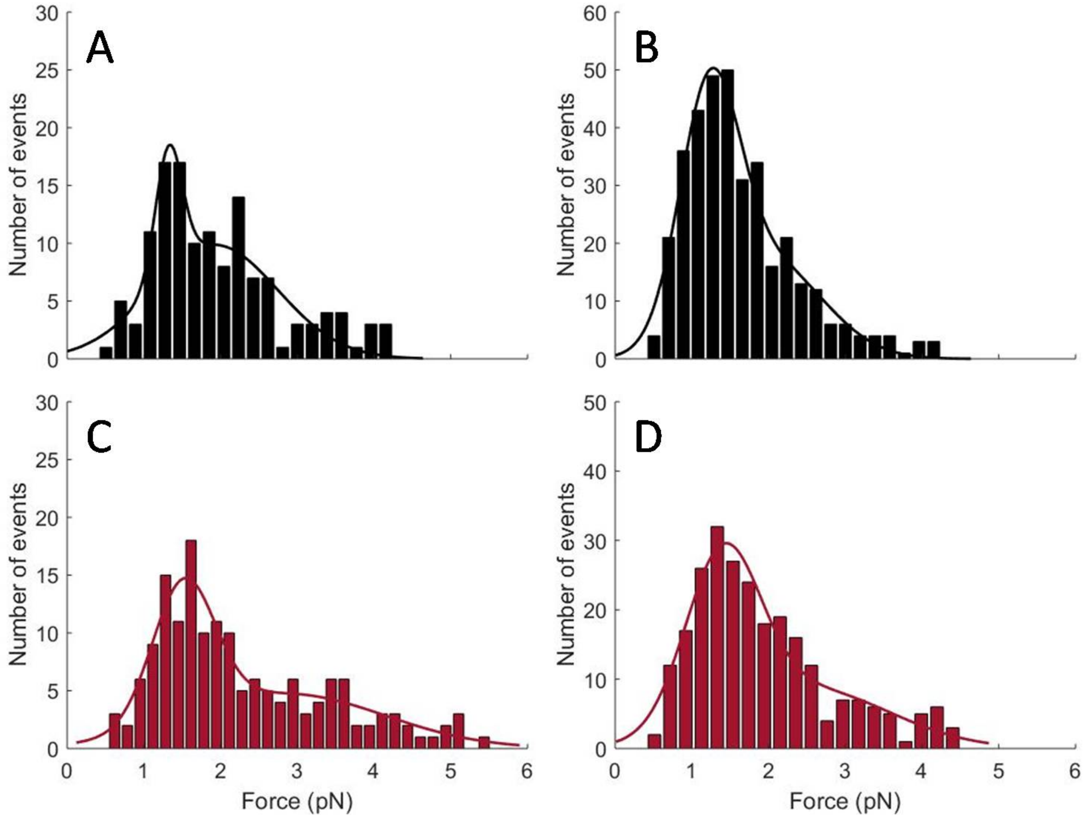


Figure S1. Intrinsic force measurements of WT and R663H human β-cardiac sS1. The data from Fig. 1B represented as force histograms. (**A**) Force histogram of a representative molecule of WT human β-cardiac sS1. (**B**) Force histogram of a representative molecule of R663H human β-cardiac sS1. (**C**) Cumulative distribution of four molecules of WT (number of events = 362). (**D**) Cumulative distribution of four molecules of R663H (number of events = 304). Each of the distributions was fitted to a double-Gaussian function and the fittings are shown.

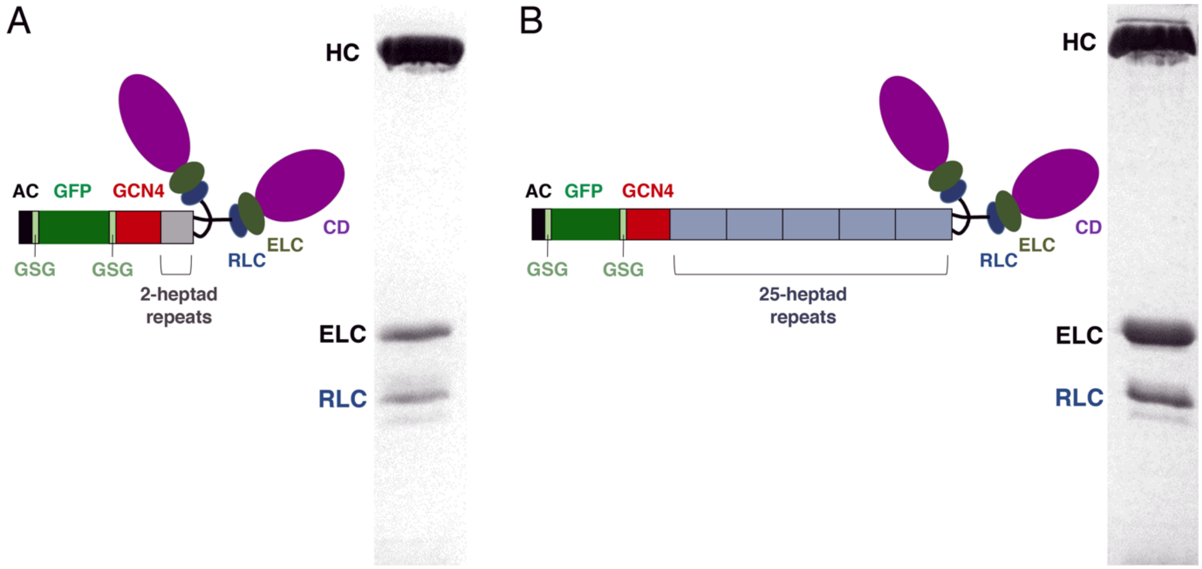


Figure S2. SDS-PAGE gel images and schematic structures of human ß-cardiac 2-hep HMM and 25-hep HMM. Each construct has 2 heavy chains, 2 ELCs and 2 RLCs. (**A**) Human ß-cardiac 2-hep HMM has 2-heptad repeats of S2 whereas (**B**) 25-hep HMM has 25-heptad repeats. The C-terminus of each construct has 1 leucine zipper (GCN4) followed by an e-GFP and an 8-residue (RGSIDTWV) PDZ-binding peptide (AC). There are flexible (GSG) linkers between GCN4 and GFP, and between GFP and AC. The gels were 15% acrylamide.

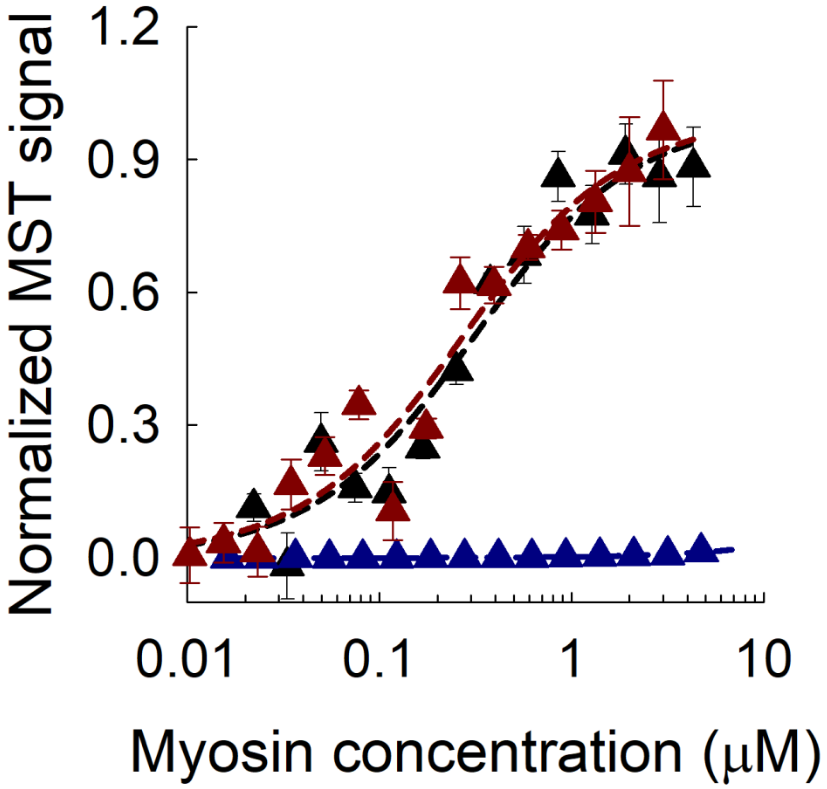


Figure S3. Binding data of human β-cardiac 25-hep HMM with the C0-C7 fragment of MyBP-C. Microscale Thermophoresis (MST) binding curves of human ß-cardiac 25-hep HMM with the C0-C7 fragment of MyBP-C for WT (black), R663H (dark red) and R403Q (blue) are shown. Experiments were performed by mixing 50 nM C0-C7 with different concentrations of 25-hep HMM. The $K_d$ values obtained from data fitting are 280 nM and 250 nM for WT and R663H, respectively. For R403Q, 25-hep HMM showed no binding to C0-C7 up to 5 μM 25-hep HMM. Data points represent the mean at each C0-C7 concentration with the s.e.m. shown. The data are the average of 3 experiments from a single set of protein preparations.
